# The BDNF Val68Met polymorphism causes a sex specific alcohol preference over social interaction and also acute tolerance to the anxiolytic effects of alcohol, a phenotype driven by malfunction of BDNF in the ventral hippocampus of male mice

**DOI:** 10.1101/2022.06.01.494180

**Authors:** Jeffrey J. Moffat, Samuel A. Sakhai, Zachary W. Hoisington, Yann Ehinger, Dorit Ron

## Abstract

The brain-derived neurotrophic factor (BDNF) Valine 66 to Methionine human polymorphism results in impaired activity-dependent BDNF release and has been linked to psychiatric disorders including depression and anxiety. We previously showed that male knock-in mice carrying the mouse Methionine homolog (Met68BDNF) exhibit excessive and compulsive alcohol drinking behaviors as compared to the wild-type Val68BDNF mice. Here, we set out to determine the potential mechanism for the heightened and compulsive alcohol drinking phenotypes detected in Met68BDNF mice. We found that male, but not female Met68BDNF mice exhibit social anxiety-like behaviors. We further show that male Met68BDNF mice exhibit a preference for alcohol over social interaction. In contrast, alcohol place preference without an alternative social reward, is similar in male Met68BDNF and Val68BDNF mice. Since the Met68BDNF mice show social anxiety phenotypes, we tested whether alcohol reliefs anxiety similarly in Met68BDNF and Val68BDNF mice and found that male, but not female Met68BDNF mice are insensitive to the acute anxiolytic action of alcohol. Finally, we show that this acute tolerance to alcohol-dependent anxiolysis can be restored by overexpressing wild-type Val68BDNF in the ventral hippocampus (vHC) of Met68BDNF mice. Together, our results suggest that excessive alcohol drinking in the Met68BDNF may be attributed, in part, to heighted social anxiety and a lack of alcohol-dependent anxiolysis, a phenotype that is associated with malfunction of BDNF signaling in the **vHC of male Met68BDNF mice.**

## Introduction

Brain-derived neurotrophic factor (BDNF) is highly expressed in the CNS and plays an important role in brain development, synaptic plasticity, learning, and memory [1]. BDNF is released both pre- and postsynaptically and its release depends on neuronal depolarization [1,2]. A single nucleotide polymorphism (SNP) within the human *BDNF* gene results in a substitution of Valine at position 66 (Val66BDNF) with Methionine (Met66BDNF), causing a reduction in the activity-dependent release of the neurotrophic factor and thus to **an attenuation of normal** BDNF-mediated signaling [3–5]. The BDNFVal66Met SNP has been linked to multiple neuropsychiatric disorders, including depression and anxiety [6,7], and several small scale studies have suggested that the BDNFVal66Met SNP is also associated with alcohol use disorder (AUD) [8–13]. In order to evaluate the impact of this BDNF SNP on alcohol related behaviors, we utilized a knock-in strategy in mice to replace the mouse Valine homolog at position 68 (Val68BDNF) with Methionine (Met68BDNF) and showed that homozygous male Met68BDNF mice consume excessive amount of alcohol [14]. We further demonstrated that the mice drink despite the addition of quinine suggesting that the Met68BDNF mice consume alcohol compulsively [14]. Finally, we reported that overexpression of Val68BDNF in the medial prefrontal cortex (mPFC) of Met68BDNF mice converts compulsive, excessive intake to moderate alcohol consumption [14].

AUD is frequently diagnosed in individuals with comorbid neuropsychiatric disorders [15–18]. For example, a global meta-analysis concluded that approximately 20-40% of individuals with major depressive disorder or anxiety also develop AUD [19]. One potential explanation for this phenomenon is the so-called “self-medication hypothesis”, which posits that individuals consume increasing quantities of alcohol in order to elicit relief from symptoms of mood disorders including social anxiety [20–23]. In fact, 10% of people suffering from AUD also endure social anxiety [23], and subjects who suffer from AUD are 4.5 times more likely to also exhibit social anxiety [24]. Heterozygous (Val/Met) and homozygous (Met/Met) genotypes have been linked with anxiety and specifically social anxiety in humans [13,25]. For instance, in analyzing the Pediatric Imaging, Neurocognition, and Genetics (PING) study [26], Li and colleagues recently uncovered a significantly higher self-reported social anxiety score in Met66BDNF allele carriers than in individuals with homozygous Val66BDNF allele [25]. The authors further showed that social deficits can be recapitulated in the mouse model of the human allele [25].

Here, we examined potential underlying causes for the heightened alcohol consumption in Met68BDNF mice, including social anxiety and alcohol-mediated anxiolysis.

## Materials and Methods

### Animals and breeding

Homozygous Val68BDNF (Val/Val) and Met68BDNF (Met/Met) generation (C57Bl6/J background) and characterization are described in Warnault et al. [14]. Female and male Val68BDNF and Met68BDNF mice were bred separately and were tested at the age of 8-10 weeks. Genotyping was performed as described previously [14]. Juvenile female and male C57Bl6/J mice (4-6 weeks) were purchased from Jackson Laboratories (Bar Harbor, Main). Mice were housed using a 12-hour light/dark cycle (lights on 7AM–7PM), with ad libitum access to food and water. All procedures were performed in accordance with guidelines from the University of California, San Francisco Institutional Care and Use Committee.

### Materials

AAV1/2-CMV-Val68BDNF-GFP (AAV-Val68BDNF; 10^12^ TU/ml) and AAV1/2-CMV-GFP (AAV-GFP; (10^12^ TU/ml) were produced by the UNC Vector Core (Chapel Hill, NC) and were characterized in Warnault et al. [14]. Ethyl alcohol (190 proof) was purchased from Thermosphere Scientific (Waltham, MA).

### Solution preparation

Alcohol solution was prepared from absolute anhydrous alcohol (190 proof) diluted to 20% alcohol (v/v) in 0.9% saline solution.

### Behavioral Assays

All behavioral analyses, with the exception of the Loss of Righting Reflex (LORR) test, were recorded and analyzed using Noldus Ethovision XT software (Wageningen, the Netherlands). Behavioral testing was performed in a dimly lit room (10-15 lux) during the light-cycle, and mice were allowed to habituate to the conditions in the room for at least one hour before testing commenced. All apparatuses were cleaned first with 70% alcohol and then with water between each animal. Test mice were habituated to experimenter handling, and intra-peritoneal (i.p.) injection by systemically administrating saline for at least three days prior to the start of experiments. Stranger mice (novel, C57Bl/6 sex matched, juvenile interaction partners used in social behavior experiments) were only used with one cohort of experimental mice and were group housed. Mice were habituated to the experimental room for at least one hour prior to beginning experiments and were habituated to wire cages, where applicable, before experiments began.

#### Three-Chamber Sociability and Social Novelty

Three-chamber sociability and social novelty tests were adapted from previous studies [27,28]. The three-chamber apparatus (40 x 60 x 25 cm) was divided into three equal zones (40 x 20 x 25 cm) by acrylic walls, connected by small doors (4 x 4 cm). Prior to the start of experiments, experimental mice were placed in the center chamber and allowed to explore the apparatus for 5 minutes before being placed in the center chambers with both side doors closed. For the sociability test, a novel, C57Bl6/J, juvenile (4-6 weeks old), sex-matched mouse (stranger I), was placed in a round wire cage (10 cm diameter) in one of the distal chambers, while an identical, empty cage was placed in the other distal chamber. The experimental mouse was then allowed to freely explore the entire apparatus for 15 minutes, after which point, the mouse was moved into the central chamber and the doors were closed. Cages were then swapped to opposite distal chambers, and a second, more-novel, C57Bl6/J, juvenile (4-6 weeks old), sex-matched mouse (stranger II) was placed inside the previously empty cage, in preparation for the social novelty test. Doors were opened, and the experimental mouse was once again allowed to freely explore the entire apparatus. During both tests, the amount of time spent in each chamber was recorded. Preference for a social partner over an empty cage in the sociability test was indicative of normal sociability. Preference for the stranger I mouse over the stranger II mouse in the social novelty task was interpreted as abnormal social behavior.

#### Open Field Social Interaction

On the day before the test, mice were habituated for 5 minutes in the open field apparatus (43 x 43 cm). Mice were then placed in the open field apparatus with a novel, wild-type, juvenile (4-6 weeks old), sex-matched interaction partner, and their interactions were recorded for 5 minutes. Cumulative body contact was calculated as time in which the center point of each animal was within 2 cm of the other. Proximity between the experimental mouse and its C57Bl6/J interaction partner was recorded within a 5 cm radius. Approach behavior was defined as vector movement in the direction of the interaction partner and retreat behavior was defined as movement away from the interaction partner.

#### Social-Alcohol Conditioned Place Preference/Aversion (CPP/CPA)

The social-alcohol CPP/CPA apparatus consisted of two large compartments (20 x 18 x 30 cm) connected by a corridor (20 x 7 x 30 cm) with one compartment consisting of lighter colored walls and mesh flooring, while the other one comprising of darker colored walls and grid rod flooring. On the first day of the experiment, mice were subjected to a pre-test during which they were allowed to freely explore the entire apparatus for 15 minutes and the time spent in each chamber was recorded. No mice reached the exclusion criteria (spend more than 70% of the time in either chamber during the pre-test). Mice were then pseudorandomly assigned to associate one chamber with social interaction and the other with alcohol (unbiased CPP/CPA). For the next 3 days, mice were conditioned to associate their social-paired chamber with social interaction and an i.p. injection of saline, and to associate the alcohol-paired chamber with an i.p. injection of alcohol (2 g/kg). Specifically, during the morning of each conditioning day, each mouse received an i.p. injection of saline before being placed in their assigned social interaction-paired chamber with a sex-matched, juvenile C57Bl/6 mouse (4-6 weeks old) for 10 minutes. In the afternoon of each conditioning day, each mouse received an i.p. injection of 20% alcohol (2 g/kg) before being placed in their assigned alcohol-paired chamber for 10 minutes. A post-test was performed on day 5, in which mice were once again allowed to freely explore the entire apparatus for 15 minutes. The time spent in each chamber during the pre-test and post-test were recorded and quantified.

#### Alcohol Conditioned Place Preference

Alcohol CPP was adapted from [29]. Specifically, the apparatus consists of two chambers (17 x 13 x 25 cm) connected by a central door. One chamber consisted of lighter colored walls and mesh flooring, while the other was made of darker colored walls and grid rod flooring. The first day of the experiment was considered a pre-test, during which test mice freely explored the entire apparatus for 15 minutes and time spent in each chamber was recorded. Using a biased design, alcohol treatment was paired with the less-preferred chamber, and the saline treatment paired with the more-preferred chamber. The following six experimental days were split into alternating days of saline or alcohol conditioning. On saline conditioning days (days 2, 4, 6), mice received an i.p. injection of saline immediately before placement in the saline-paired chamber for 5 minutes. On alcohol conditioning days (days 3, 5, 7), mice received an i.p. injection of alcohol (2 g/kg) before being placed in the alcohol-paired chamber for 5 minutes. Day 8 of the experiment consisted of a post-test, during which test mice freely explored the apparatus for 15 minutes. The time spent in each chamber during the pre-test and post-test were recorded and quantified.

#### Loss of Righting Reflex

LORR was conducted as described previously [30]. Mice received an i.p. injection of a hypnotic dose of alcohol (4.0 g/kg) before being placed in a clean plexiglass cage. The amount of time it took for each animal to lose its’ righting reflex (latency) was recorded. At this point, mice were returned to their home cage and placed on their backs. The amount of time it took for each mouse to regain its’ righting reflex, e.g., the time that it took mice to right themselves from their backs 3 times in one minute (duration) was measured.

#### Elevated Plus Maze

Elevated Plus Maze (EPM) assay was adapted from [31]. Specifically, mice were injected with either saline or alcohol (1.25 g/kg) and 10 minutes later were placed on the central platform (5 x 5 cm) of an apparatus elevated 40 cm above the floor, facing one of two closed arms (30 x 5 x 15 cm), which are perpendicular to two open arms (30 x 5 cm). Mice were allowed to freely explore the apparatus for 5 minutes, and the total amount of time each mouse spent exploring the open arms, and the distal portion of both open arms (outer 15 cm), was recorded and quantified.

### Stereotaxic surgery

Met68BDNF mice (8-10 weeks old) underwent stereotaxic surgery as described in [14,32] targeting the vHC (Franklin and Paxinos stereotaxic atlas, 3rd edition). Specifically, mice were anesthetized by vaporized isoflurane, and were placed in a digital stereotaxic frame (David Kopf Instruments, Tujunga, CA). Two holes were drilled above the site of the injection and the injectors (stainless tubing, 33 gauges; Small Parts Incorporated, Logansport, IN) were then slowly lowered into the vHC (AP: −3.0, ML: ±3.0, DV: −3.55, infusion at −3.5 from bregma). The injectors were connected to Hamilton syringes (10 μl; 1701; Harvard Apparatus, Holliston, MA), and infusion of 1 μl of virus was controlled by an automatic pump at a rate of 0.1 μl/min (Harvard Apparatus, Holliston, MA). The injectors remained in place for an additional 10 minutes to allow the virus to diffuse and were then gently removed. Mice were allowed to recover in their home cages for at least 3 weeks before further testing to allow for maximal overexpression of Val68BDNF [14].

### Confirmation of viral expression

Characterization of infection and overexpression was performed as described previously [14,32]. At the end of experiments animals were euthanized by cervical dislocation, and the brains were removed. Brain regions were isolated from a 1-mm-thick coronal section dissected on ice, and a green fluorescence (GFP) signal was visualized and recorded using an EVOS FL tabletop fluorescent microscope (ThermoFisher Scientific; Waltham, MA).

### Data analysis

Graphpad Prism 9 was used for statistical analysis. D’Agostino–Pearson normality test was used to verify the normal distribution of variables. Data were analyzed using two- or three-way ANOVA, with or without repeated measures, and student’s t-test, where appropriate. For two- and three-way ANOVAs, significant main effects or interactions were calculated using Šidák’s multiple comparisons test. p value cutoff for statistical significance was set to 0.05.

## Results

### Aberrant social behavior in male but not female Met68BDNF mice

Male Met68BDNF mice consume alcohol excessively and compulsively [14], however, the underlying cause of this phenotype is unknown. Humans carrying the Met66BDNF allele show increased social anxiety [25], and stress and anxiety including social anxiety are thought to be major contributors to AUD [19,23]. Thus, to determine if the mouse BDNFVal68Met polymorphism increases susceptibility for social anxiety in female and male mice, we first measured sociability and social novelty behaviors using a three-chamber social interaction paradigm [27,28]. In the sociability test phase, mice freely explored an apparatus containing, in one distal chamber, a novel, juvenile, sex-matched, C57Bl/6J mouse, identified as “stranger I”. The other distal chamber contained an empty cage (Figure 1A. left). Next, in the social novelty test phase, the chambers were swapped, and a new social interaction partner identified as “stranger II”, was placed inside the previously empty cage (Figure 1A, right), and interaction time was recorded. We found that female and male Val68BDNF and Met68BDNF mice spent a similar amount of time in a chamber containing the stranger I mouse (Figure 1B-F, Two-way ANOVA; interaction effect, F(1,53) = 1.495, p = 0.2268; main effect of genotype, F (1, 53) = 0.8987, p = 0.3474; main effect of sex, F (1, 53) = 58.83, p < 0.0001) suggesting that sociability *per se* is not affected by the SNP. However, we did detect a significant sex difference between male and female mice (Figure 1B). Mice from all test groups spent more time in a chamber containing a stranger I mouse than they did in an empty chamber (Figure 1C, Three-way ANOVA; main effect of chamber, F(1, 53) = 44.90, p < 0.0001; main effect of genotype, F(1, 53) = 0.0001945, p = 0.9889; main effect sex, F(1, 53) = 169.4, p < 0.0001; chamber x genotype x sex, F(1, 53) = 2.259, p = 0.1388; Male Val68BDNF Empty vs. Male Val68BDNF Stranger I: p = 0.04, Male Met68BDNF Empty vs. Male Met68BDNF Stranger I: p < 0.0001, Female Val68BDNF Empty vs. Female Val68BDNF Stranger I: p = 0.0186, Female Met68BDNF Empty vs. Female Met68BDNF Stranger I: p = 0.05). In the social novelty paradigm in which mice could choose between spending time with the familiar mouse (strange I) or a novel mouse (stranger II), Female Val68BDNF and Met68BDNF spent an equal amount of time interacting with stranger II mice and female Met68BDNF mice spent significantly more time interacting with the stranger II mouse than they did with the stranger I mouse (Figure 1 D-F; Two-way ANOVA, interaction effect, F (1, 54) = 16.57, p = 0.0002, main effect of sex, F (1, 54) = 7.237, p = 0.0095, main effect of genotype, F (1, 54) = 5.633, p = 0.0212; Female Val68BDNF vs. Female Met68BDNF: p = 0.403, Male Val68BDNF vs. Male Met68BDNF: p <0.0001). Male Val68BDNF mice spent an equal amount of time interacting with “stranger I” and “stranger II” mouse (Figure 1E-F, Three-way ANOVA; main effect of chamber, F(1, 53) = 0.4522, p = 0.5042; main effect of genotype, F(1, 53) = 0.1209, p = 0.7294; main effect sex, F(1, 53) = 155.3, p < 0.0001; chamber x genotype, F(1, 53) = 6.852, p = 0.0115; chamber x genotype x sex, F(1, 53) = 11.35, p = 0.0014; Male Val68BDNF Stranger I vs. Male Val68BDNF Stranger II: p = 0.5585, Male Met68BDNF Stranger I vs. Male Met68BDNF Stranger II: p < 0.0001, Female Val68BDNF Stranger I vs. Female Val68BDNF Stranger II: p = 0.2204, Female Met68BDNF Stranger I vs. Female Met68BDNF Stranger II: p = 0.0315). In contrast, male Met68BDNF mice spent significantly less time interacting with the “stranger II” mouse as compared to their wild-type Val68BDNF counterparts (Figure 1D, F) and significantly more time interacting with the stranger I mouse than the stranger II mouse (Figure 1E-F). These findings suggest that male Met68BDNF exhibit social anxiety-like behavior or a diminished rewarding sensation due to social novelty.

**Figure 1.**
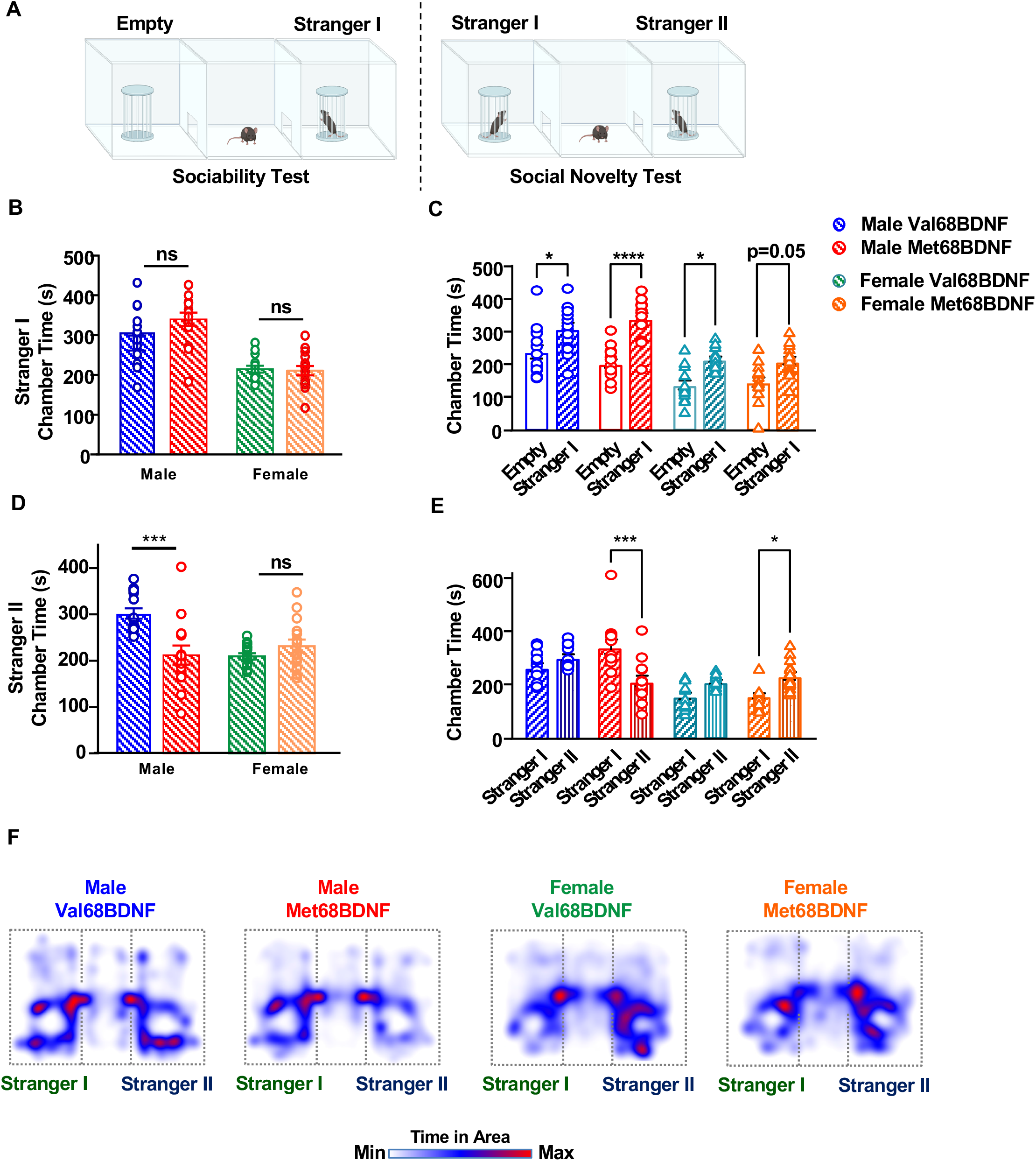
Male Met68BDNF mice exhibit social anxiety-like behaviors. (**A**) *Three-chamber sociability and social novelty paradigms. Sociability test* (left). The distal chamber contained an empty cage, and the other chamber contained a sex match juvenile C57Bl/6J mouse (Stranger I). Female (green) and male (blue) Val68BDNF, and female (orange) and male (red) Met68BDNF mice were allowed to interact with the Stranger I mouse for 15 minutes, and the time a mouse spent with Stranger I mouse was recorded and quantified. *Social novelty test* (right). Stranger I was relocated to the distal chamber, and a sex matched novel juvenile C57Bl/6J mouse (Stranger II), was placed in the other chamber. Female (green) and male (blue), Val68BDNF and female (orange) and male (red) Met68BDNF mice were allowed to interact with stranger I or Stranger II for 15 minutes, and the time a mouse spent with stranger I or stranger II was recorded and quantified. (**B**) Female and male Val68BDNF and Met68BDNF mice spent about the same amount of time interacting with the stranger I mouse during the sociability test. (**C**) Male Met68BDNF mice and male and female Val68BDNF mice spent significantly more time in the chamber containing the stranger I mouse than they do in an empty chamber. (**D**) Male Met68BDNF mice spent significantly less time interacting with a stranger II mouse in the social novelty test as compared to male Val68BDNF mice. Female Val68BDNF and Met68BDNF mice spent a similar amount of time interacting with a stranger II mouse. (E) Male Met68BDNF mice spent significantly more time in a chamber containing the more-familiar stranger I than they do in a chamber containing stranger II. Female Met 68BDNF mice spent significantly more time in the chamber containing stranger II than the chamber containing stranger I. (**F**) Heatmaps of mean mouse position for male and female Val68BDNF and Met68BDNF mice during the social novelty test shown in (C, E). Stranger I mice are shown on the left of each heatmap, and stranger II are shown on the right side. Dotted lines indicate the approximate position of apparatus walls. All data are represented as mean ± SEM. *** p < 0.001, * p < 0.05, ns = nonsignificant. Female Val68BDNF: n = 15, all other experimental groups: n=14.

To further explore the extent of aberrant social behavior in male Met68BDNF mice, female and male Val68BDNF and Met68BDNF mice were subjected to an open field social interaction paradigm. In this assay, test mice were paired with a novel, juvenile, sex-matched, C57Bl/6J partner in an empty open field apparatus (Figure 2A). Male Met68BDNF mice spent significantly less time in physical contact with a novel juvenile male mouse than male Val68BDNF mice (Figure 2B, Two-tailed Student’s t-test, p < 0.01, t=2.947, df=17). Additional analysis of male Met68BDNF mice during open field social interaction did not reveal any one particular aspect of aberrant social interaction, however, as nose-to-nose, nose-to-tail, approach, and retreat interactions trended lower than those seen in male Val68BDNF mice but were not statistically significant (Supplemental Figure 1, Unpaired t-test, t=1.589, df=17, p = 0.1305; 1B, Unpaired t-test, t=1.608, df=17, p = 0.1263; 1C, Unpaired t-test, t=1.345, df=17, p = 0.1964; 1D, Unpaired t-test, t=0.9802, df=17, p = 0.3407; 1E, Unpaired t-test, t=1.504, df=17, p = 0.1508; 1F, Unpaired t-test, t=1.596, df=17, p = 0.1289; 1G, Unpaired t-test, t=1.304, df=17, p = 0.2098; 1H, Unpaired t-test, t=1.018, df=17, p = 0.3228; 1I, Unpaired t-test, t=1.294, df=17, p = 0.2131; 1J, Unpaired t-test, t=0.8372, df=17, p = 0.4141). In contrast, female Val68BDNF and Met68BDNF mice spent equal amounts of time in physical contact with a juvenile, female, C57Bl/6J mouse (Figure 2C, Two-tailed Student’s t-test, t=0.5858, df=26, p = 0.5631). Together, these data unravel a sex-specific, genotype-dependent impairment in social behaviors in Met68BDNF mice suggestive of social anxiety.

**Figure 2.**
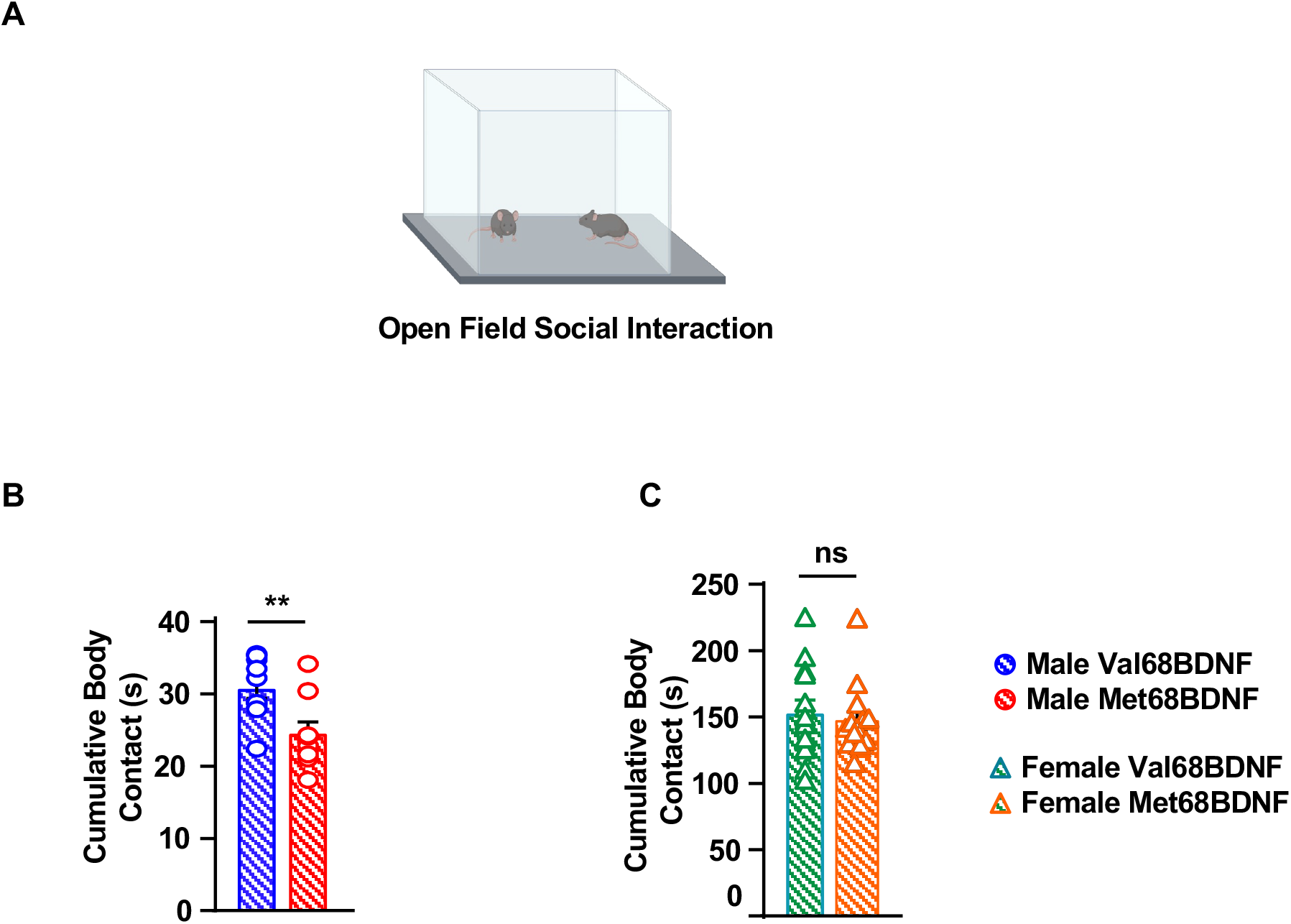
Male Met68BDNF mice exhibit social interaction anxiety-like behavior. (**A**) *Open field social interaction paradigm*. Male (blue) and female (green) Val68BDNF and male (red) and female (orange) Met68BDNF mice were placed in an open field apparatus with a novel sex matched juvenile C57Bl/6J mouse for 10 minutes and body contacts were recorded. (**B**) Male Met68BDNF mice spent significantly less time in contact with a novel interaction partner than male Val68BDNF. (**C**) There was no significant difference in total body contacts with a novel interaction partner between female Val68BDNF and Met68BDNF mice. ** p < 0.01, ns = non-significant. (**B**): Male Val68BDNF: n = 10, Male Met68BDNF: n = 9, (**C**): Female Val68BDNF and Met68BDNF n = 14.

### Male Met68BDNF mice exhibit social aversion and alcohol preference in a social-alcohol conditioned place preference/aversion test

Social anxiety is a risk factor for AUD [23,24], and previous research demonstrated that humans carrying at least one Met66BDNF allele report higher levels of social anxiety [25]. Since male Met68BDNF mice exhibit a social anxiety phenotype, we examined the possibility that mice carrying the mutation prefer alcohol over social interaction. To test this possibility, we conducted a social-alcohol conditioned place preference/conditioned place aversion (CPP/CPA) paradigm, in which mice were conditioned to associate one distinct chamber with social interaction with a novel male juvenile partner each morning for 3 days, and a second distinct chamber with an acute alcohol administration (2g/kg) each afternoon for 3 days (Figure 3A). We then compared the time each mouse spent freely exploring each chamber following conditioning (post-test) with the amount of time mice spent exploring the same chambers prior to conditioning (pre-test). We found that male Val68BDNF mice found alcohol and social interaction equally rewarding, as evidenced by the amount of time mice spent exploring the social-paired and alcohol-paired chambers in the post-test, compared with during the pre-test (Figure 3B-C, Three-way ANOVA; main effect of conditioning, F(1, 20) = 7.826, p = 0.0111; main effect of reward, F(1, 20) = 0.1998, p = 0.6596; main effect of genotype, F(1,20) = 0.01176, p = 0.9147; conditioning x reward x genotype. F(1, 20) = 7.076, p=0.0150; Pre/social Val68BDNF vs. Post/social Val68BDNF: p = 0.2406, Pre/alcohol Val68BDNF vs. Post/alcohol Val68BDNF: p = 0.9528, Pre/social Met68BDNF vs. Post/social Met68BDNF: p = 0.0002, Pre/alcohol Met68BDNF vs. Post/alcohol Met68BDNF: p = 0.0028). In contrast, male Met68BDNF mice exhibited a significantly lower preference for the social interaction chamber and a higher preference for the alcohol-paired chamber (Figure 3B-C), indicative of a simultaneous social aversion and alcohol preference, potentially due either to a devaluation of social reward, a hypervaluation of alcohol reward or both.

**Figure 3.**
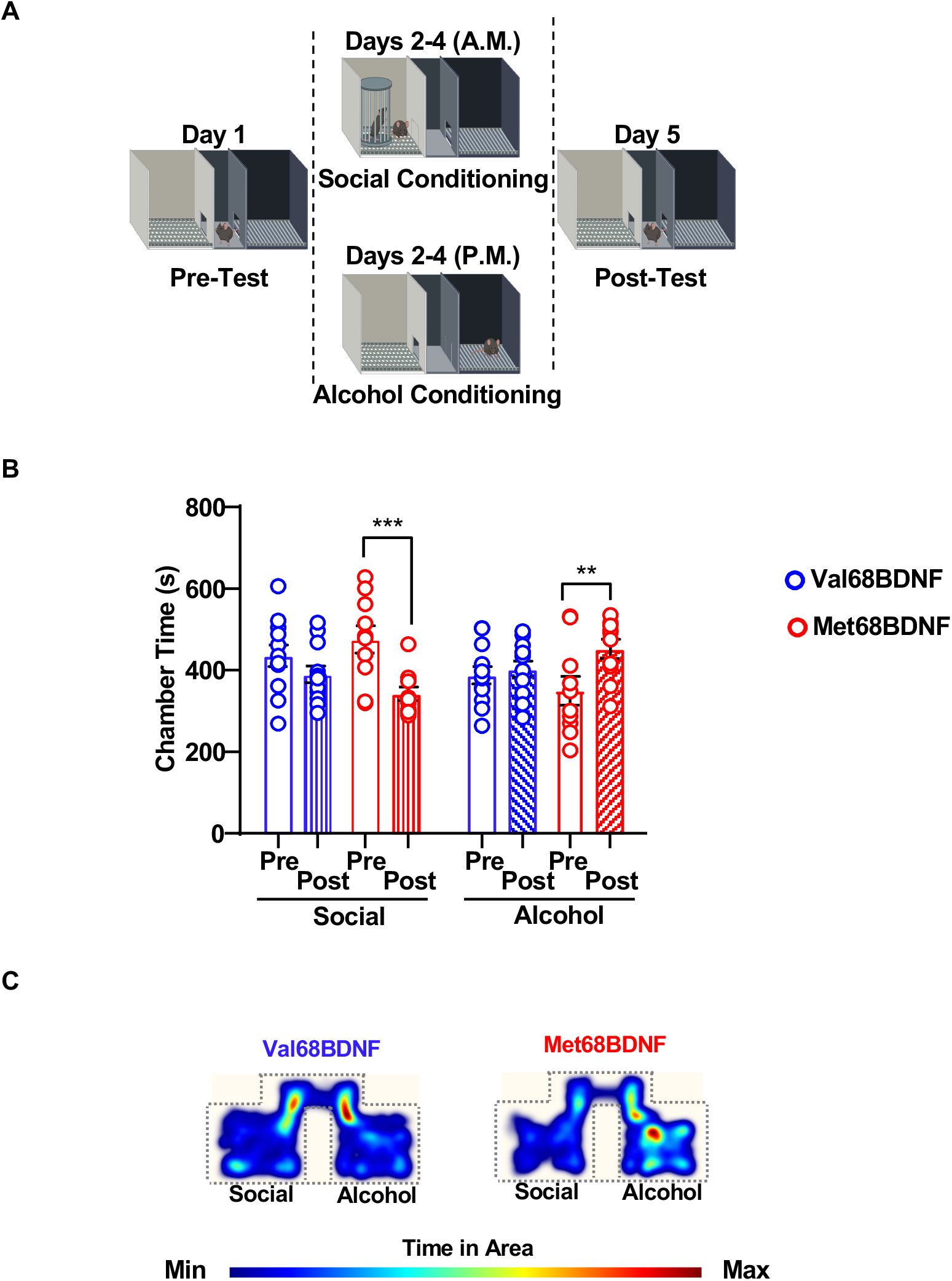
Male Met68BDNF mice demonstrate social aversion and alcohol preference in a social-alcohol place conditioning test. (**A**) *Outline of the social-alcohol place preference/aversion test*. On day 1, mice explored the entire apparatus for 15 minutes. In the morning of days 2-4 conditioning days, mice received an i.p. injection of saline before a 10-minute social interaction period in the social assigned chamber. In the afternoon of days 2-4 mice received an i.p. injection of 2 g/kg of alcohol prior to being placed in the alcohol-paired chamber for 10 minutes. On day 5, mice were allowed to freely explore the entire apparatus for 15 minutes and the time spend in the social and alcohol chambers were recorded and quantified. (**B**) Male Met68BDNF mice (red) spent significantly less time in the social-paired chamber in the post-test than they did in the pre-test, and significantly more time in the alcohol-paired chamber in the post-test compared to the pre-test. Male Val68BDNF mice (blue) do not exhibit place preference or aversion. (**C**) Representative heatmaps of mouse position for Val68BDNF and Met68BDNF cohorts during the social-alcohol CPP/CPA shown in (**B**). The social chamber is depicted on the left and the alcohol chamber is depicted on the right for each heatmap. The upper portion of the heatmap represents the connecting hallway between chambers. Dotted lines indicate the approximate position of apparatus walls. All data are represented as mean ± SEM; ** p < 0.01, *** p < 0.001. Val68BDNF: n = 12, Met68BDNF: n = 10.

Next, to determine whether male Met68BDNF mice prefer the alcohol chamber over social interaction because they find alcohol more rewarding, we used an alcohol CPP paradigm (Supplemental Figure 2A). In the post-test, male Val68BDNF and Met68BDNF mice spent significantly more time in the alcohol-paired chamber following conditioning and less time in the saline-paired chamber in the posttest, compared to the pre-test (Supplemental Figure 2B, Two-way ANOVA; interaction effect, F(1,24) = 0.09766, p = 0.7574; main effect of genotype, F (1, 24) = 1.052, p = 0.3153, main effect of conditioning, F (1, 24) = 19.68, p = 0.0002; Pre Val68BDNF vs. Post Val68BDNF: p = 0.0144, Pre Met68BDNF vs. Post Met68BDNF: p = 0.0036). These results demonstrate that Met68BDNF mice experience the rewarding effects of alcohol to a similar degree as Val68BDNF mice, suggesting that Met68BDNF mice prefer the alcohol-associated chamber because of social anxiety and/or devalued social rewards, and not because of amplified alcohol rewards.

### Male Met68BDNF mice are resistant to alcohol-induced sedation

Alcohol tolerance has been associated with increased propensity to develop AUD [33–35]. We therefore examined the possibility that the BDNFVal68/Met polymorphism alters male mice sensitivity to the acute actions of alcohol. To explore this possibility, we tested whether male Met68BDNF mice exhibit an abnormal response to an acute sedative dose of alcohol as compared to male Val68BDNF mice. Specifically, we utilized a loss of righting reflex (LORR) paradigm and measured how long it took mice to lose their righting reflex after receiving a systemic administration of a hypnotic dose (4g/kg) of alcohol, and how long the LORR lasted in each animal. We found that it took male Met68BDNF mice significantly longer to lose their righting reflex as compared to male Val68BDNF mice (Supplemental Figure 3A, Two-tailed Student’s t-test, t = 2.439, df =13, p < 0.05). Furthermore, following LORR onset, male Met68BDNF mice also exhibited a significantly quicker recovery from the sedative effects of alcohol, as indicated by a group mean LORR duration being less than half as long as the mean duration for male Val68BDNF mice (Supplemental Figure 3B, Two-tailed Student’s t-test, t = 3.736, df = 13, p < 0.01). Together, these results suggest that male Met68BDNF exhibit tolerance to the acute effects of a hypnotic dose of alcohol.

### Male Met68BDNF mice exhibit acute tolerance to the anxiolytic action of alcohol

Next, we examined whether acute alcohol tolerance is more generalized and also whether alcohol is potentially perceived differently in the two genotypes. To do so, we evaluated the acute anxiolytic properties of alcohol in male Met68BDNF mice as compared to male Val68BDNF. We also determined the anxiolytic action of alcohol in female Val68BDNF and Met68BDNF mice. To do so, we used an elevated plus maze paradigm (EPM) in which mice received a systemic dose of alcohol (1.25 g/kg) or saline and 10 minutes later were placed in the center of an elevated plus maze apparatus, which is composed of two enclosed arms and two open arms that do not have barriers on their perimeters (Figure 4A). Administration of alcohol (1.25 g/kg) significantly increased the time mice spent in the open arms (Figure 4B-F, Three-way ANOVA; main effect of alcohol treatment, F (1, 75) = 64.07, p < 0.0001, main effect of sex, F (1,75) = 21.56, p < 0.0001, main effect of genotype, F (1, 75) = 0.9477, p = 0.3334, treatment x sex, F(1, 75) = 2.391, p = 0.1263, treatment x genotype, F(1, 75) = 0.09304, p = 0.7612), and specifically in the distal portion of the open arms compared with mice receiving saline, suggesting an anxiolytic response to alcohol (Figure 4C-F, Three-way ANOVA; main effect of alcohol treatment, F (1, 75) = 44.25, p < 0.0001, main effect of sex, F (1,75) = 36.33, p < 0.0001, main effect of genotype, F (1, 75) = 0.00053, p = 0.9817, treatment x sex, F(1, 75) = 7.033, p = 0.0098, treatment x genotype, F(1, 75) = 0.2250, p = 0.6366; Saline Male Val68BDNF vs. Alcohol Male Val68BDNF: p = 0.0051). We found a significant difference of the effect of alcohol between male and female mice (Figure 4B-C, F). We also observed a phenotypic difference in the EPM that was not captured in the statistical analysis of open arm time due to differences in baseline levels between the male Val68BDNF and Me68BDNF mice. Therefore, we performed an additional analysis in which we normalized the time in open arms after alcohol treatment to the time in open arm of the saline group (Figure 4D-E). We found that male Met68BDNF mice exhibit a significant resistance to the anxiolytic effects of alcohol, as acute systemic administration of 1.25 g/kg of alcohol did not significantly alter male Met68BDNF mice’s exploration time in the open arm (Figure 4D; Two-way ANOVA interaction effect, F(1, 52) = 8.233, p = 0.0059, main effect of alcohol treatment, F (1, 52) = 35.17, p < 0.0001; main effect of genotype, F (1, 52) = 8.233, p = 0.0059; Saline Male Val68BDNF vs. Alcohol Male Val68BDNF: p < 0.0001, Saline Male Met68BDNF vs. .Alcohol Male Met68BDNF: p = 0.075) or distal open arm (Figure 4E; Two-way ANOVA interaction effect, F(1, 52) = 8.651, p = 0.0049, main effect of alcohol treatment, F (1, 52) = 17.94, p < 0.0001; main effect of genotype, F (1, 52) = 8.651, p = 0.0049; Saline Male Val68BDNF vs. Alcohol Male Val68BDNF: p < 0.0001, Saline Male Met68BDNF vs. Alcohol Male Met68BDNF: p = 0.6064) as compared to male saline-treated male Met68BDNF mice. The difference in the male Met68BDNF mice was not due to changes in locomotion in response to alcohol administration (Supplemental Figure 4). Overall, these data demonstrate a sexspecific, genotype-dependent impairment in alcohol-mediated anxiolysis (Figure 4). Our data further show that male mice carrying the Met68BDNF allele exhibit tolerance to the acute actions of alcohol.

**Figure 4.**
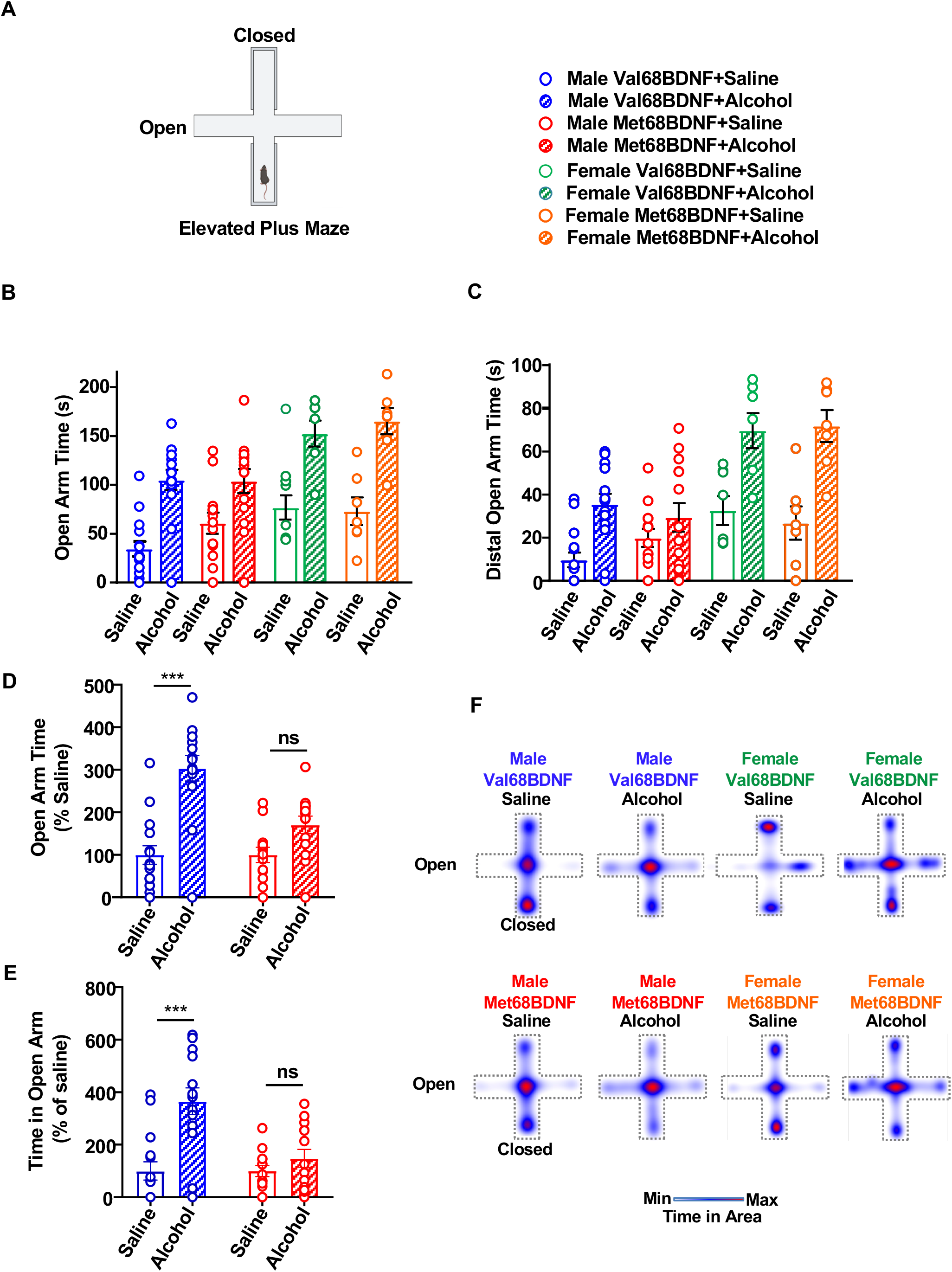
Male Met68BDNF exhibit acute tolerance to the anxiolytic action of alcohol. (**A**) Ten minutes following i.p. injection of saline or alcohol (1.25 g/kg), mice were placed in the center of the elevated plus maze and allowed to freely explore the apparatus. The time mice spent in the open arm and in the distal portion of the open arms was measured. (**B-C**) Following an i.p. administration of alcohol (hatched bars), female and male mice spent significantly more time in the open arm (**B**) and the distal portion of the open arm (**C**) than upon receiving an injection of saline (open bars). Male Met68BDNF mice (red) exploration time in the open arm (**B**) and the distal portion of the open arm (**C**) was unaltered following a systemic administration of alcohol (hatched bars) or saline (empty bars). (**D-E**) Additional analysis on male mice was performed in which the time in open arms or distal arms after alcohol treatment was normalized to the time in open arm or distal arms of the saline group. (**F**) Heatmaps representing mean relative position of animals in each group. Open arms are represented horizontally, and closed arms extend vertically from the center. Dotted lines indicate the approximate position of apparatus walls/ledges. Data are represented as mean ± SEM; * p < 0.05, ** p < 0.01,*** p < 0.001. Male Val68BDNF + saline: n = 15, male Val68BDNF + alcohol: n = 14, male Met68BDNF + saline: n = 13, male Met68BDNF + alcohol: n = 14, female Val68BDNF + saline: n = 6, female Val68BDNF + alcohol: n = 7, female Met68BDNF + saline: n = 7, female Met68BDNF + alcohol: n = 7.

### Overexpression of Val68BDNF in the ventral hippocampus of male Met68BDNF mice restores the anxiolytic effects of alcohol

The ventral hippocampus (vHC) plays a role in anxiety-like behaviors in rodents [36–41], but also with anxiolysis [42]. We therefore hypothesized that BDNF in the vHC plays a role in dampening anxiety-like behaviors and in promoting alcohol-dependent anxiolysis. We further hypothesized that malfunction of BDNF signaling in the vHC manifests in behaviors such as resistance to the anxiolytic actions of alcohol. To test this possibility, the vHC of male Met68BDNF mice was infected with adeno-associated virus (AAV) expressing either wild-type Val68BDNF or a GFP control (Figure 5A). Three weeks later, a timepoint in which viral infection is maximal, mice received systemic administration of saline or alcohol (1.25 g/kg) 10 minutes prior to placement on the EPM apparatus. Similar to results reported in Figure 4, male Met68BDNF infected with AAV-GFP in the vHC were resilient to the anxiolytic effects of alcohol and spent a similar amount of time in the open arms (Figure 5B,D, Two-way ANOVA; interaction effect, F(1, 38) = 5.644, p = 0.0227; main effect of alcohol treatment, F(1, 38) = 28.34, p < 0.0001; main effect of virus, F(1, 38) = 2.964, p = 0.0933; AAV-GFP Saline vs. AAV-GFP Alcohol: p = 0.0769, AAV-BDNF Saline vs. AAV-BDNF Alcohol: p < 0.0001), and in their distal portions following saline or alcohol administration (Figure 5C, D, Two-way ANOVA, interaction effect, F(1, 38) = 3.204, p = 0.0814, main effect of alcohol treatment, F (1, 38) = 31.08, p < 0.0001; main effect of virus, F (1, 38) = 4.126, p = 0.0493; AAV-GFP Saline vs. AAV-GFP Alcohol: p = 0.0542, AAV-BDNF Saline vs. AAV-BDNF Alcohol: p < 0.0001). In contrast, male Met68BDNF mice infected with Val68BDNF in the vHC spent significantly more time in the open arms and distal open arms following alcohol administration (Figure 5B-D). Importantly the rescue of alcohol-dependent anxiolysis was not due to alterations in locomotion (Supplemental Figure 5, Two-way ANOVA; interaction effect, F(1, 38) = 0.03114, p = 0.8609; main effect of alcohol treatment, F(1, 38) = 10.64, p = 0.0023; main effect of virus, F(1, 38) = 0.2184, p = 0.643). Together, these data imply that deficits in BDNF/TrkB signaling in vHC circuitry in carriers of the Met68BDNF allele contribute to impaired alcohol-mediated anxiolysis and to the development of acute tolerance.

**Figure 5.**
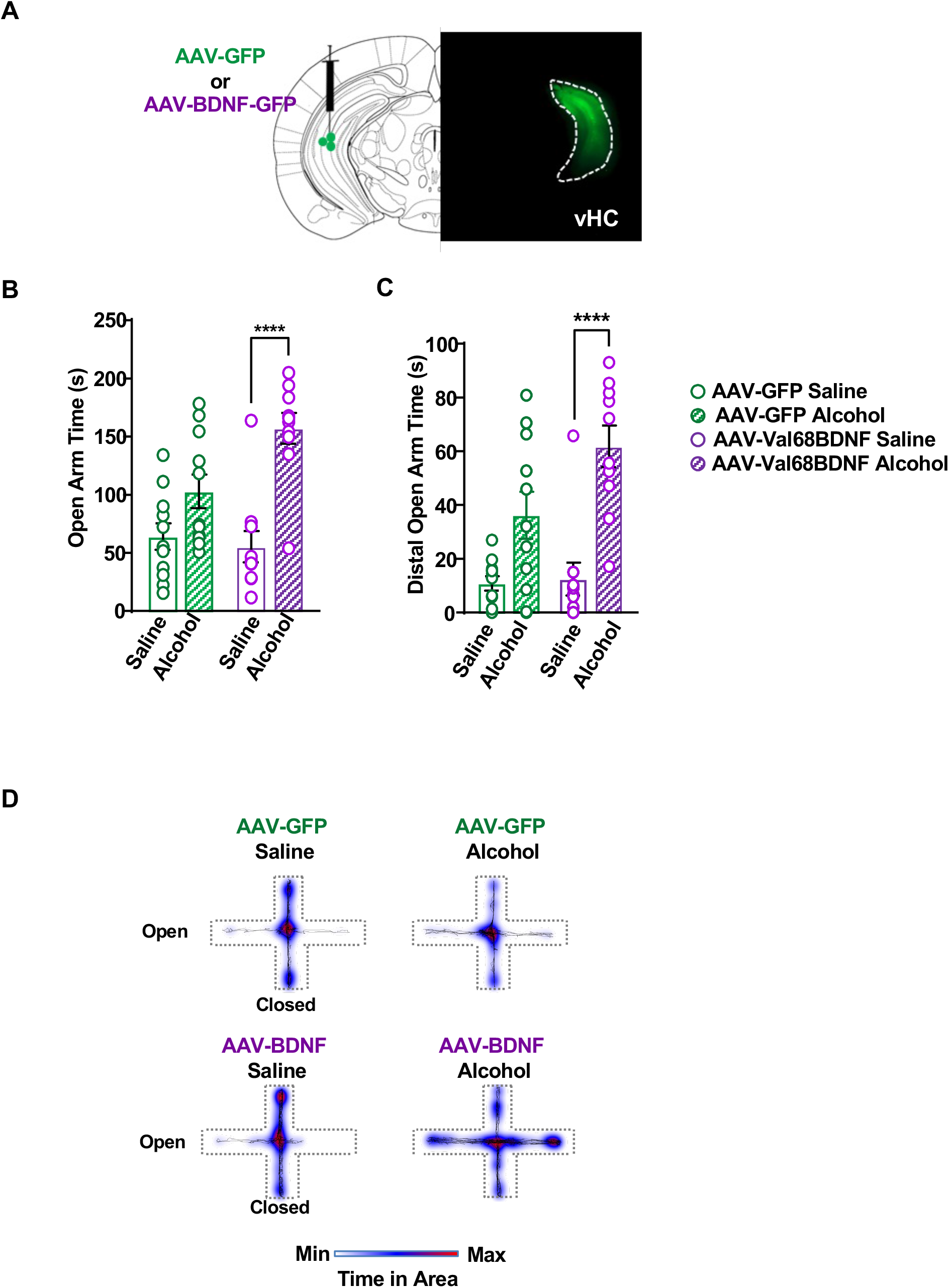
Overexpressing BDNF in the vHC of male Met68BDNF rescues the anxiolytic effect of alcohol. (**A**) Confirmation of Val68BDNF overexpression. Image (10x) depicts GFP signal in the vHC of a Met68BDNF mouse that received AAV-Val68BDNF in the vHC. (B-C). Male Met68BDNF that received AAV-GFP (green) in the vHC spent similar time in both the open arm (**B**) and the distal open arm (C) following i.p. injection of alcohol (1.25 g/kg) (hatched bars) or saline (open bars). Met68BDNF mice that received AAV-Val68BDNF (purple) in the vHC spent significantly more time in the open arm (**B**) and the distal open arm (C) following a systemic administration of alcohol (hatched bars) than they did following saline administration (open bars). (**D**) Heatmaps of group mean position on the elevated plus maze. Open arms are horizontal, and closed arms extend vertically from the center. Dotted lines indicate the approximate position of apparatus walls/ledges. Data represented as mean ± SEM; *** p < 0.001. Male Met68BDNF + AAV-GFP: n = 11, Male Met68BDNF + AAV-Val68BDNF: n = 10.

## Discussion

In this study we provide evidence to suggest that male carriers of the Met68BDNF allele exhibit social anxiety, alcohol preference over social interaction and acute tolerance to the sedative and anxiolytic actions of alcohol. We further show that overexpression of the wild-type Val68BDNF in the vHC of the Met68BDNF mice restores normal anxiolytic responses to alcohol suggesting that proper BDNF signaling in the vHC is required for alcohol-dependent anxiolysis.

Li et al., showed that male carriers of the human or mouse Met66BDNF allele show social anxiety phenotypes [25], and we obtained similar data in male mice carrying the Met68BDNF allele. Li et al. suggested that the social deficits in the male Met66BDNF mice are caused by reduced function of BDNF in a medial orbitofrontal cortical (mOFC) to basolateral amygdala (BLA) circuit during development [25]. Li et al. further showed that these deficits can only be rescued when wild-type Val66BDNF is overexpressed during a peri-adolescent developmental window of Met66BDNF mice but not during adulthood [25]. Although we did not specifically examine whether social anxiety can be rescued in the adult Met68BDNF mice, we were able to rescue compulsive alcohol drinking phenotype by overexpressing Val68BDNF in the mPFC of adult Met68BDNF mice [14], and here we show that alcohol-dependent anxiolysis can be restored by overexpressing Val68BDNF in the vHP of adult Met68BDNF mice. These data suggest that malfunction of BDNF during development does not play a role in all tested behaviors. Another important difference between the two mouse lines is that we did not observe a baseline anxiety differences in mice carrying the Val68BDNF and Met68BDNF alleles ([14] and data herein), whereas mice generated by Li and colleagues carrying the human Met66BDNF allele, show heighten basal anxiety levels [43]. Furthermore, Vandenberg et al. compared the behaviors of Met66BDNF and Met68BDNF mice as well as their wild-type counterparts in a battery of cognitive paradigms and found numerous additional differences between the two mouse lines [44]. It is important to note that our knockin mouse line and Li and colleagues knockin mouse line are different. For example, Chen et. al kept a Histidine tail at the C-terminal of the BDNF protein [43], whereas our mouse line did not contain additional amino acids [14]. This difference is of importance as a Histidine tag can produce **non-**physiological changes to the function of the endogenous protein. First, the addition of multiple Histidines was shown to increase protein stability [45], and also to change the confirmation and activity of recombinant proteins [46,47]. Thus, differences in Met66BDNF and Met68BDNF half-life and/or confirmation could play a role in the behavior of the mouse. It is also plausible that because one or all of these differences Chen et al. mice [43] have developmental deficits [25] whereas ours do not. More studies are required to clarify why mice carrying the val66/68MetBDNF alleles show such differences in neurobehavioral phenotypes.

We found that male Met68BDNF exhibited aversion to a social-paired chamber and a preference for an alcohol-paired chamber in a social-alcohol CPP/CPA test. However, alcohol-place preference was normal in Met68BDNF mice suggesting that social deficits and not increased alcohol reward are the primary driving factor of mice preferring the chamber that was associated with alcohol and not the chamber that was associated with social interaction. Ten percent of subjects suffering from AUD also endure social anxiety [48], and those who suffer from AUD are 4.5 times more likely to also exhibit social anxiety [23,24,48,49]. In addition, social anxiety disorders are strongly correlated with increased risk for heavy alcohol use [48–51]. Furthermore, social stress during adolescence increases alcohol intake in male and female C57Bl/6 mice [52], and predicts alcohol intake in humans [53]. Together, these reports show that social anxiety is a risk factor for heavy alcohol use. We previously showed that male Met68BDNF mice consume alcohol both excessively and compulsively [14]. Several human studies report higher average alcohol consumption in Met66BDNF allele carriers [12,13]. Thus, it is plausible that subjects carrying the BDNF SNP consume alcohol excessively in part to alleviate social anxiety. Therefore, it would be of great interest to determine whether social anxiety and AUD are correlated in human carriers of the Met66BDNF allele.

We observed that male Met68BDNF mice are resistant to the acute sedative and anxiolytic actions of alcohol suggesting that this mutation leads to the development of acute alcohol tolerance. It is unlikely, that the acute tolerance to is due to enhanced alcohol metabolism for two reasons: First, blood alcohol concentration was the same in Val68BDNF and Met68BDNF mice 90 minutes after receiving a dose of 2.5g/kg of alcohol [14], and second, since overexpression of wild-type Val68BDNF in the vHC of Met68BDNF mice was sufficient to rescue alcohol-dependent anxiolysis. Acute tolerance is a well-known characteristic of AUD [35], and classic longitudinal studies by Schuckit and colleagues showed that heightened acute alcohol tolerance, coincides with an increased risk for AUD [33,34]. Not much is known about the mechanisms underlying alcohol tolerance [35]. Thus, acute tolerance as shown herein, and compulsive alcohol consumption as shown in [14] may be linked and are due to a malfunction of a single gene and its downstream signaling.

Sex differences in social behavior have been widely described [54–59], and we observed sex differences in social behaviors between the male and female Met68BDNF mice. Specifically, disruption in BDNF function does not seem to affect the social behavior phenotypes in female Met68BDNF, and in contrast male Met68BDNF mice exhibit social anxiety. In line with our findings, human data indicates for example that male, but not female Met66BDNF allele carriers experience higher incidences of major depressive disorder [60]. We also observed that, whereas male Met68BDNF mice are resistant to the anxiolytic action of alcohol, female Met68BDNF mice are not, as they show a similar pattern of behavior to Val68BDNF female and male mice. These data suggest that sex differences may be more generalized in mice carrying the Met68BDNF allele. Interestingly, although we did not compare them directly, female Val68BDNF and Met68BDNF mice exhibited divergent open field social interaction behavioral phenotypes, compared to their male counterparts. Because this sex difference exists in both genotypes, it is unlikely to be related to BDNF. However, further studies are needed to determine whether BDNF in female mice plays a role in other behaviors, including gating alcohol use, which is mediated in part via corticostriatal circuitries in male mice [14,61–63].

We establish that BDNF in vHC neurons, plays a critical role in mediating alcohol-induced anxiolysis. BDNF acts both in an autocrine and a paracrine manner. Specifically, BDNF is released in an activity dependent manner postsynaptically, and more commonly through axonal terminals [1,2]. Thus, it is plausible that BDNF synthesized in the vHC is released postsynaptically and activates TrkB receptors in dendrites of the same neurons or presynaptically by targeting other neurons within the vHP. Alternatively, and more likely, BDNF produced in vHC neurons and released in target regions may influence behaviors via signaling in those brain regions. For example, the vHC extends neuronal projections to, and receives them from, the BLA and the mPFC, and connections between these three brain regions are linked with anxiety and specifically with social anxiety-like behaviors in rodents [37,42,64–66]. Furthermore, vHC neurons projecting to the lateral septum (LS) were shown to suppress anxiety-like behavior [42] whereas vHC to mPFC circuits promote anxiety [40,42]. In contrast, vHC neurons that project to the nucleus accumbens (NAc) [67] or mPFC [68] were shown to drive social reward memory. In rats, alcohol dependence has been reported to specifically increase synaptic excitability in the vHC [69], and inactivation of a projection from the ventral subiculum of the hippocampus to the NAc shell decreases context-induced alcohol relapse [70]. Moreover, we previously found that overexpressing wildtype BDNF in the mPFC of Met68BDNF mice is sufficient to reverse compulsive alcohol consumption in adult mice [14] which raises the possibility that overexpression of wild-type BDNF in the mPFC mimics the endogenous BDNF in vHC to mPFC circuit. Future studies are required to map vHC BDNF neurons and their target regions and to determine whether BDNF in these circuits plays a role in other alcohol-related phenotypes including social anxiety phenotypes, alcohol preferences vs. social interaction, and compulsive alcohol drinking.

In summary, in this study we demonstrate that male Met68BDNF mice exhibit social anxiety-like phenotypes and are resistant to the acute behavioral effects of alcohol. The present study and Warnault et al. [14] suggest that BDNF is crucially involved in gating excessive and compulsive alcohol intake [14], in dampening social deficits including social aversion in the context of alcohol place preference as well as in promoting alcohol anxiolysis. However, whether these behavioral phenotypes overlap is as of yet unknown and need to be further explored. Our data also bring forward the importance of BDNF-expressing neurons in the vHC in the anxiolytic actions of alcohol, and additional research is needed to identify the mechanism by which BDNF regulates the behavior.

## Funding and Disclosure

This research was supported by the National Institute of Alcohol Abuse and Alcoholism R37AA01684 (DR) and F32AA028422 (JJM). None of the authors have a conflict of interest.

## Acknowledgments

The authors thank JiHwan Yu for his help. Some of the figures were made in part using BioRender.

## Author Contributions

JJM contributed to experimental design, data acquisition, data analysis, and manuscript preparation. SAS contributed to experimental design and data acquisition. ZWH contributed to data acquisition. YE contributed to data analysis and manuscript preparation. DR conceived the study. contributed to experimental design and manuscript preparation.

## Supplemental Material

**Supplemental Figure 1.**
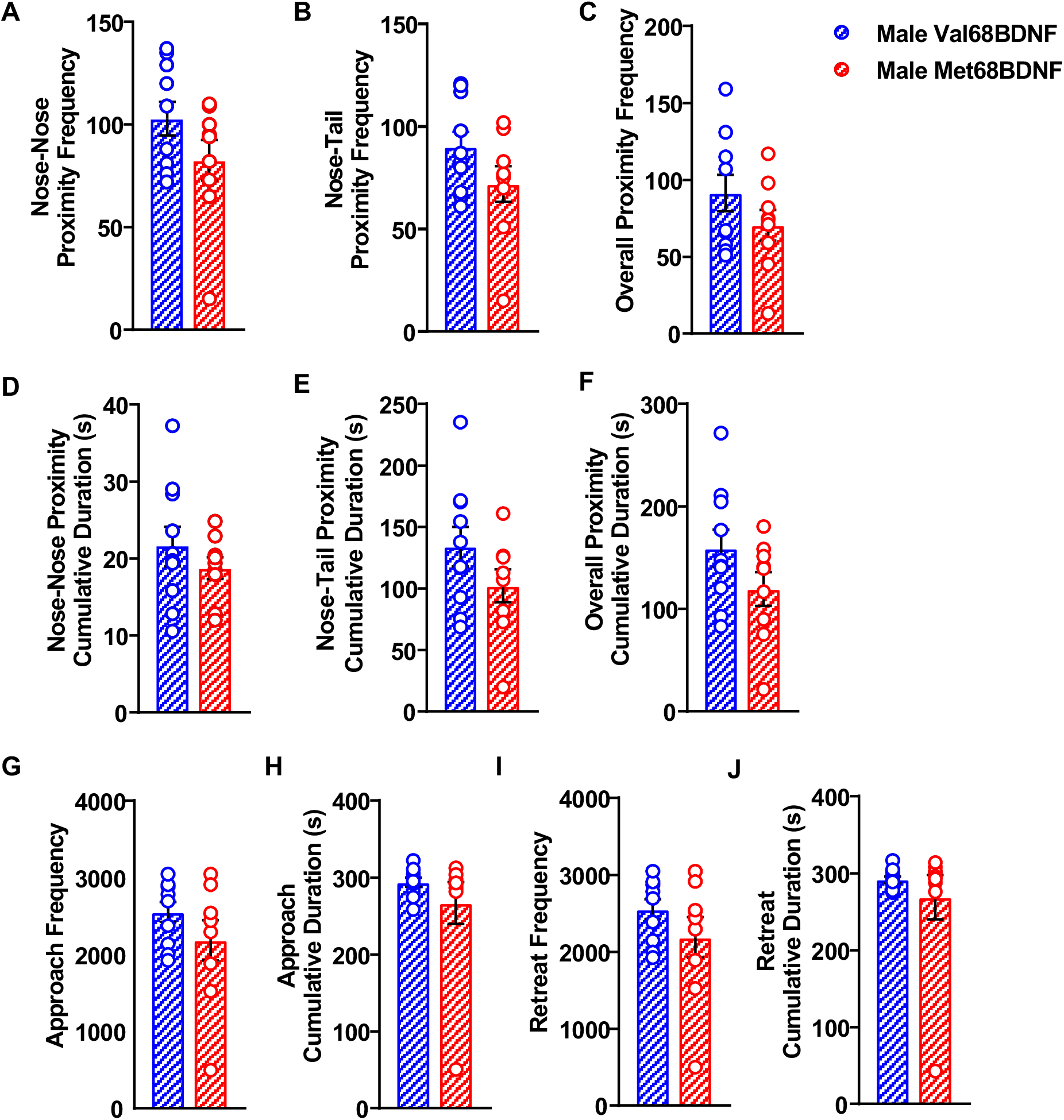
Male Met68BDNF mice do not exhibit any specific deficits in open field social interaction beyond a general reduction in interaction. (**A**) Male Val68BDNF and Met68BDNF mice came within 5 cm of their interaction partner (nose-to-nose) a similar number of times in the open field social interaction test (Unpaired t-test, t=1.589, df=17, p = 0.1305). (**B**) Male Val68BDNF and Met68BDNF mice came within 5 cm of their interaction partner (nose-to-tail) a similar number of times in the open field social interaction test (Unpaired t-test, t=1.608, df=17, p = 0.1263). (**C**) Male Val68BDNF and Met68BDNF mice came within 5 cm of their interaction partner a similar number of times in the open field social interaction test overall (Unpaired t-test, t=1.345, df=17, p = 0.1964). (**D**) Male Val68BDNF and Met68BDNF mice spent a similar amount of time within 5 cm of their interaction partner (nose-to-nose) in the open field social interaction test (Unpaired t-test, t=0.9802, df=17, p = 0.3407). (**E**) Male Val68BDNF and Met68BDNF mice spent a similar amount of time within 5 cm of their interaction partner (nose-to-tail) in the open field social interaction test (Unpaired t-test, t=1.504, df=17, p = 0.1508). (**F**) Male Val68BDNF and Met68BDNF mice spent a similar amount of time within 5 cm of their interaction partner in the open field social interaction test overall (Unpaired t-test, t=1.596, df=17, p = 0.1289). (**G**) Male Val68BDNF and Met68BDNF mice approached their interaction partner a similar number of times in the open field social interaction test (Unpaired t-test, t=1.304, df=17, p = 0.2098). (**H**) Male Val68BDNF and Met68BDNF mice spent a similar amount of time approaching their interaction partner in the open field social interaction test (Unpaired t-test, t=1.018, df=17, p = 0.3228). (**I**) Male Val68BDNF and Met68BDNF mice retreated from their interaction partner a similar number of times in the open field social interaction test (Unpaired t-test, t=1.294, df=17, p = 0.2131). (**J**) Male Val68BDNF and Met68BDNF mice spent a similar amount of time retreating from their interaction partner in the open field social interaction test (Unpaired t-test, t=0.8372, df=17, p = 0.4141). Data are represented as mean ± SEM. Val68BDNF: n = 10, Met68BDNF: n = 9.

**Supplemental Figure 2.**
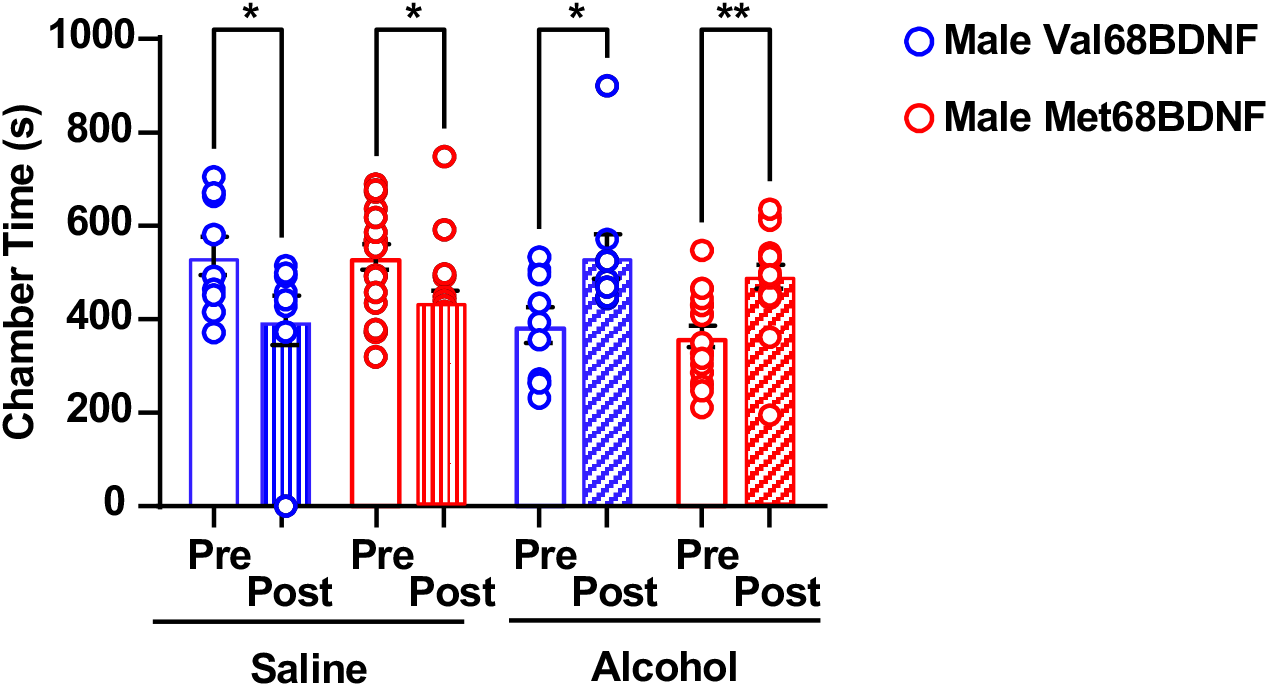
Male Val68BDNF and Met68BDNF mice exhibit alcohol place preference. Mice underwent an alcohol place preference paradigm in which they were first able to freely explore both chambers (pre-test). On alternating conditioning days, mice were placed in a saline- or alcohol-paired chamber after receiving an i.p. injection of saline or 2 g/kg of alcohol. During the post-test day, mice were once again allowed to freely explore the entire apparatus and time spend in each of the chambers were recorded and quantified. Male Val68BDNF (blue) and Met68BDNF (red) mice exhibit a significantly higher preference for the alcohol-paired chamber compared with the saline-paired chamber (**Two-way ANOVA, p < 0.05, main effect of alcohol, F (1, 24) = 14.94, p < 0.001)**. All data are represented as mean ± SEM; * p < 0.05. Val68BDNF: n = 9 (2 were removed due to health issues), Met68BDNF: n = 17.

**Supplemental Figure 3.**
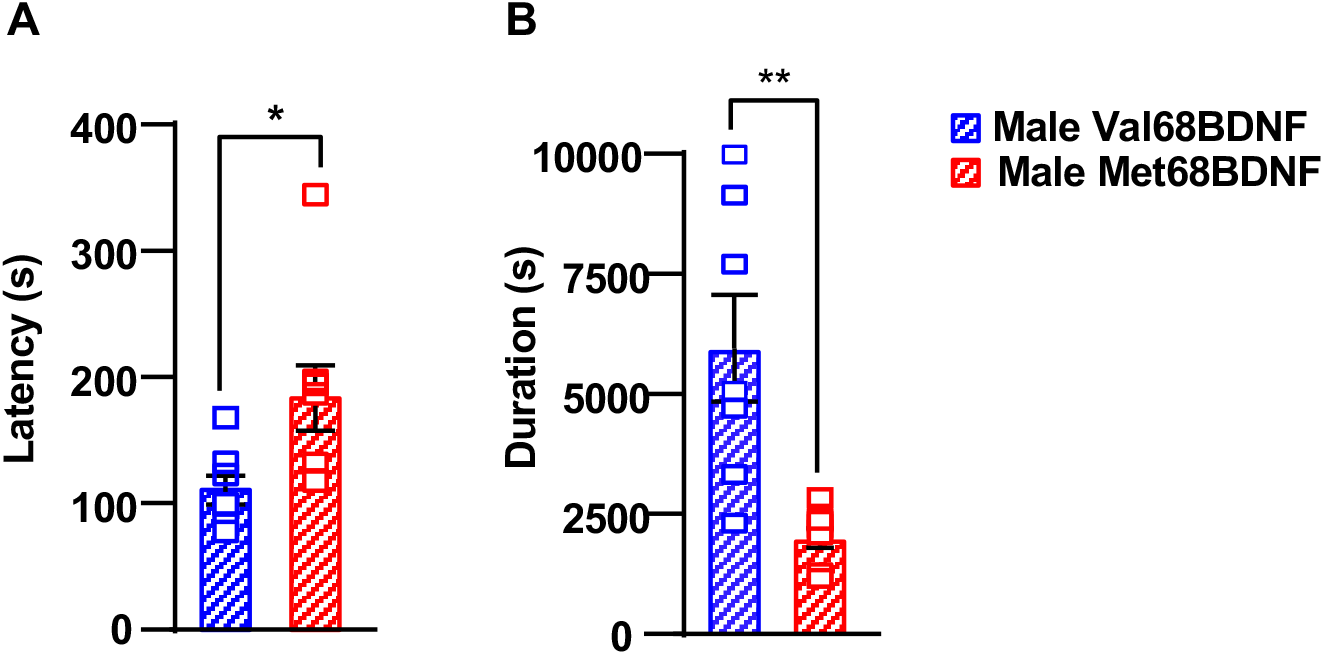
Male Met68BDNF mice are resistant to the sedative effects of alcohol. (**A**) Mice received 4 g/kg of alcohol and the time it took for sedation to set in, and the duration of the sedation were recorded. The latency between alcohol injection (4 g/kg) and the point at which mice do not right themselves after being placed on their backs was significantly greater in male Met68BDNF mice (red) than male Val68BDNF mice (blue) **(Two-tailed Student’s t-test, p < 0.05, t = 2.439, df = 13**). (**B**) The total duration of LORR for male Met68BDNF mice (red) was significantly shorter than it was for mice Val68BDNF mice (blue) (**Two-tailed Student’s t-test, p < 0.01, t = 3.736, df = 13**). Data are represented as mean ± SEM; * p < 0.05, ** p < 0.01. Val68BDNF: n = 7, Met68BDNF: n = 8.

**Supplemental Figure 4.**
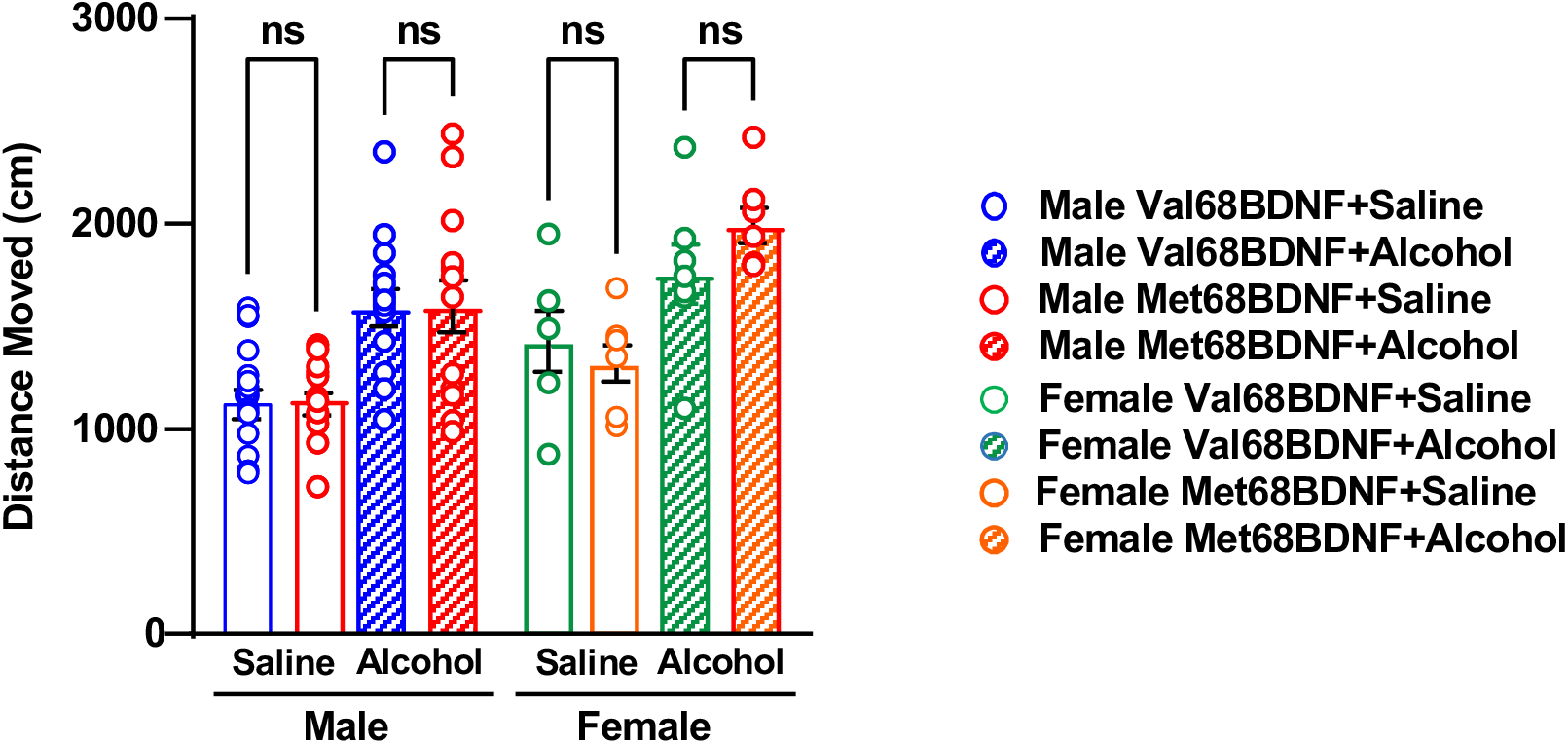
No genotype differences in male and female Val68BDNF and Met68BDNF locomotion in the elevated plus maze. Male Met68BDNF (red) mice travel the same distance on the elevated plus maze as male Val68BDNF (blue) mice following i.p. injection of saline (empty bars) or 1.25 g/kg of alcohol (hashed bars). Female Val68BDNF (green) and Met68BDNF (orange) mice also travel the same distance after i.p. injection of saline (empty bars) or 1.25 g/kg of alcohol (hatched bars) (**Two-way ANOVA, main effect of alcohol treatment, F (3, 75) = 13.34, p < 0.0001; main effect of sex (1,75) = 10.93, p < 0.01**). Data are represented as mean ± SEM. Male Val68BDNF + saline: n = 15, male Val68BDNF + alcohol: n = 14, male Met68BDNF + saline: n = 13, male Met68BDNF + alcohol: n = 14, female Val68BDNF + saline: n = 6, female Val68BDNF + alcohol: n = 7, female Met68BDNF + saline: n = 7, female Met68BDNF + alcohol: n = 7.

**Supplemental Figure 5.**
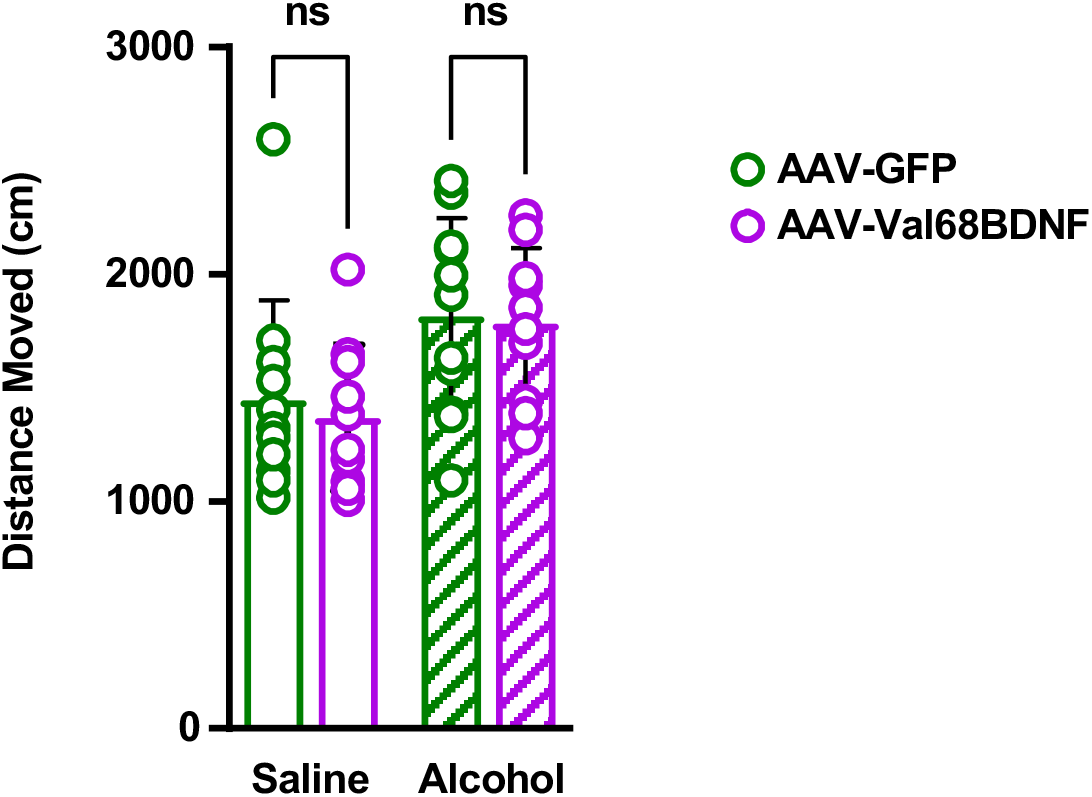
BDNF overexpression in the vHC of male Met68BDNF mice does not impact alcohol-dependent locomotion in the elevated plus maze. **The BDNF Val68Met polymorphism causes a sex specific alcohol preference over social interaction and also acute tolerance to the anxiolytic effects of alcohol, a phenotype driven by malfunction of BDNF in the ventral hippocampus of male mice** Male Met68BDNF mice that received AAV-Val68BDNF in the vHC (purple) move the same distance as male Met68BDNF mice that received AAV-GFP in the vHC (green) following i.p.injection of either saline (empty bars) or 1.25 g/kg of alcohol (hatched bars) (Two-way ANOVA, main effect of alcohol treatment, F (1, 38) = 10.64, p < 0.01). Data represented as mean ± SEM. Male Met68BDNF + AAV-GFP: n = 11, Male Met68BDNF + AAV-Val68BDNF: n = 10.

## Notes

### Competing Interest Statement

The authors have declared no competing interest.

### Summary of Updates

Figure 4 was updated

## References

1 Wang CS, Kavalali ET, Monteggia LM. BDNF signaling in context: From synaptic regulation to psychiatric disorders. Cell. 2022;185(1):62–76.

2 De Vincenti AP, Rios AS, Paratcha G, Ledda F. Mechanisms That Modulate and Diversify BDNF Functions: Implications for Hippocampal Synaptic Plasticity. Front Cell Neurosci. 2019;13:135.

3 Egan MF, Kojima M, Callicott JH, Goldberg TE, Kolachana BS, Bertolino A, et al. The BDNF val66met polymorphism affects activity-dependent secretion of BDNF and human memory and hippocampal function. Cell. 2003;112(2):257–69.

4 Chen ZY, Patel PD, Sant G, Meng CX, Teng KK, Hempstead BL, et al. Variant brain-derived neurotrophic factor (BDNF) (Met66) alters the intracellular trafficking and activity-dependent secretion of wild-type BDNF in neurosecretory cells and cortical neurons. J Neurosci. 2004;24(18):4401–11.

5 Chen ZY, Ieraci A, Teng H, Dall H, Meng CX, Herrera DG, et al. Sortilin controls intracellular sorting of brain-derived neurotrophic factor to the regulated secretory pathway. J Neurosci. 2005;25(26):6156–66.

6 Dincheva I, Glatt CE, Lee FS. Impact of the BDNF Val66Met polymorphism on cognition: implications for behavioral genetics. Neuroscientist. 2012;18(5):439–51.

7 Notaras M, Hill R, van den Buuse M. The BDNF gene Val66Met polymorphism as a modifier of psychiatric disorder susceptibility: progress and controversy. Mol Psychiatry. 2015;20(8):916–30.

8 Benzerouk F, Gierski F, Gorwood P, Ramoz N, Stefaniak N, Hubsch B, et al. Brain-derived neurotrophic factor (BDNF) Val66Met polymorphism and its implication in executive functions in adult offspring of alcohol-dependent probands. Alcohol. 2013;47(4):271–4.

9 Grzywacz A, Samochowiec A, Ciechanowicz A, Samochowiec J. Family-based study of brain-derived neurotrophic factor (BDNF) gene polymorphism in alcohol dependence. Pharmacol Rep. 2010;62(5):938–41.

10 Matsushita S, Kimura M, Miyakawa T, Yoshino A, Murayama M, Masaki T, et al. Association study of brain-derived neurotrophic factor gene polymorphism and alcoholism. Alcohol Clin Exp Res. 2004;28(11):1609–12.

11 Nees F, Witt SH, Dinu-Biringer R, Lourdusamy A, Tzschoppe J, Vollstadt-Klein S, et al. BDNF Val66Met and reward-related brain function in adolescents: role for early alcohol consumption. Alcohol. 2015;49(2):103–10.

12 Wojnar M, Brower KJ, Strobbe S, Ilgen M, Matsumoto H, Nowosad I, et al. Association between Val66Met brain-derived neurotrophic factor (BDNF) gene polymorphism and post-treatment relapse in alcohol dependence. Alcohol Clin Exp Res. 2009;33(4):693–702.

13 Colzato LS, Van der Does AJ, Kouwenhoven C, Elzinga BM, Hommel B. BDNF Val66Met polymorphism is associated with higher anticipatory cortisol stress response, anxiety, and alcohol consumption in healthy adults. Psychoneuroendocrinology. 2011;36(10):1562–9.

14 Warnault V, Darcq E, Morisot N, Phamluong K, Wilbrecht L, Massa SM, et al. The BDNF Valine 68 to Methionine Polymorphism Increases Compulsive Alcohol Drinking in Mice That Is Reversed by Tropomyosin Receptor Kinase B Activation. Biol Psychiatry. 2016;79(6):463–73.

15 Melchior M, Prokofyeva E, Younes N, Surkan PJ, Martins SS. Treatment for illegal drug use disorders: the role of comorbid mood and anxiety disorders. BMC Psychiatry. 2014;14:89.

16 Sorensen T, Jespersen HSR, Vinberg M, Becker U, Ekholm O, Fink-Jensen A. Substance use among Danish psychiatric patients: a cross-sectional study. Nord J Psychiatry. 2018;72(2): 130–36.

17 Preuss UW, Gouzoulis-Mayfrank E, Havemann-Reinecke U, Schafer I, Beutel M, Hoch E, et al. Psychiatric comorbidity in alcohol use disorders: results from the German S3 guidelines. Eur Arch Psychiatry Clin Neurosci. 2018;268(3):219–29.

18 Walters RK, Polimanti R, Johnson EC, McClintick JN, Adams MJ, Adkins AE, et al. Transancestral GWAS of alcohol dependence reveals common genetic underpinnings with psychiatric disorders. Nat Neurosci. 2018;21(12):1656–69.

19 Castillo-Carniglia A, Keyes KM, Hasin DS, Cerda M. Psychiatric comorbidities in alcohol use disorder. Lancet Psychiatry. 2019;6(12):1068–80.

20 Khantzian EJ. The self-medication hypothesis of addictive disorders: focus on heroin and cocaine dependence. Am J Psychiatry. 1985;142(11):1259–64.

21 Khantzian EJ. The self-medication hypothesis of substance use disorders: a reconsideration and recent applications. Harv Rev Psychiatry. 1997;4(5):231–44.

22 Turner S, Mota N, Bolton J, Sareen J. Self-medication with alcohol or drugs for mood and anxiety disorders: A narrative review of the epidemiological literature. Depress Anxiety. 2018;35(9):851–60.

23 Bowen MT, George O, Muskiewicz DE, Hall FS. Factors Contributing to the Escalation of Alcohol Consumption. Neurosci Biobehav Rev. 2022;132:730–56.

24 Buckner JD, Turner RJ. Social anxiety disorder as a risk factor for alcohol use disorders: a prospective examination of parental and peer influences. Drug Alcohol Depend. 2009;100(1-2):128–37.

25 Li A, Jing D, Dellarco DV, Hall BS, Yang R, Heilberg RT, et al. Role of BDNF in the development of an OFC-amygdala circuit regulating sociability in mouse and human. Mol Psychiatry. 2021;26(3):955–73.

26 Jernigan TL, Brown TT, Hagler DJ, Jr., Akshoomoff N, Bartsch H, Newman E, et al. The Pediatric Imaging, Neurocognition, and Genetics (PING) Data Repository. Neuroimage. 2016;124(Pt B):1149–54.

27 Moy SS, Nadler JJ, Perez A, Barbaro RP, Johns JM, Magnuson TR, et al. Sociability and preference for social novelty in five inbred strains: an approach to assess autistic-like behavior in mice. Genes Brain Behav. 2004;3(5):287–302.

28 Kaidanovich-Beilin O, Lipina T, Vukobradovic I, Roder J, Woodgett JR. Assessment of social interaction behaviors. J Vis Exp. 2011(48).

29 Laguesse S, Morisot N, Shin JH, Liu F, Adrover MF, Sakhai SA, et al. Prosapip1-Dependent Synaptic Adaptations in the Nucleus Accumbens Drive Alcohol Intake, Seeking, and Reward. Neuron. 2017;96(1): 145–59 e8.

30 Yaka R, Tang KC, Camarini R, Janak PH, Ron D. Fyn kinase and NR2B-containing NMDA receptors regulate acute ethanol sensitivity but not ethanol intake or conditioned reward. Alcohol Clin Exp Res. 2003;27(11):1736–42.

31 Walf AA, Frye CA. The use of the elevated plus maze as an assay of anxiety-related behavior in rodents. Nat Protoc. 2007;2(2):322–8.

32 Ehinger Y, Morisot N, Phamluong K, Sakhai SA, Soneja D, Adrover MF, et al. cAMP-Fyn signaling in the dorsomedial striatum direct pathway drives excessive alcohol use. Neuropsychopharmacology. 2021;46(2):334–42.

33 Schuckit MA. Ethanol-induced changes in body sway in men at high alcoholism risk. Arch Gen Psychiatry. 1985;42(4):375–9.

34 Schuckit MA, Smith TL, Clarke D, Mendoza LA, Kawamura M, Schoen L. Predictors of Increases in Alcohol Problems and Alcohol Use Disorders in Offspring in the San Diego Prospective Study. Alcohol Clin Exp Res. 2019;43(10):2232–41.

35 Elvig SK, McGinn MA, Smith C, Arends MA, Koob GF, Vendruscolo LF. Tolerance to alcohol: A critical yet understudied factor in alcohol addiction. Pharmacol Biochem Behav. 2021;204:173155.

36 Ciocchi S, Passecker J, Malagon-Vina H, Mikus N, Klausberger T. Brain computation. Selective information routing by ventral hippocampal CA1 projection neurons. Science. 2015;348(6234):560–3.

37 Felix-Ortiz AC, Tye KM. Amygdala inputs to the ventral hippocampus bidirectionally modulate social behavior. J Neurosci. 2014;34(2):586–95.

38 Jimenez JC, Su K, Goldberg AR, Luna VM, Biane JS, Ordek G, et al. Anxiety Cells in a Hippocampal-Hypothalamic Circuit. Neuron. 2018;97(3):670–83 e6.

39 Kjelstrup KG, Tuvnes FA, Steffenach HA, Murison R, Moser EI, Moser MB. Reduced fear expression after lesions of the ventral hippocampus. Proc Natl Acad Sci U S A. 2002;99(16):10825–30.

40 Padilla-Coreano N, Bolkan SS, Pierce GM, Blackman DR, Hardin WD, Garcia-Garcia AL, et al. Direct Ventral Hippocampal-Prefrontal Input Is Required for Anxiety-Related Neural Activity and Behavior. Neuron. 2016;89(4):857–66.

41 Kheirbek MA, Drew LJ, Burghardt NS, Costantini DO, Tannenholz L, Ahmari SE, et al. Differential control of learning and anxiety along the dorsoventral axis of the dentate gyrus. Neuron. 2013;77(5):955–68.

42 Parfitt GM, Nguyen R, Bang JY, Aqrabawi AJ, Tran MM, Seo DK, et al. Bidirectional Control of Anxiety-Related Behaviors in Mice: Role of Inputs Arising from the Ventral Hippocampus to the Lateral Septum and Medial Prefrontal Cortex. Neuropsychopharmacology. 2017;42(8):1715–28.

43 Chen ZY, Jing D, Bath KG, Ieraci A, Khan T, Siao CJ, et al. Genetic variant BDNF (Val66Met) polymorphism alters anxiety-related behavior. Science. 2006;314(5796):140–3.

44 Vandenberg A, Lin WC, Tai LH, Ron D, Wilbrecht L. Mice engineered to mimic a common Val66Met polymorphism in the BDNF gene show greater sensitivity to reversal in environmental contingencies. Dev Cogn Neurosci. 2018;34:34–41.

45 Khan F, Legler PM, Mease RM, Duncan EH, Bergmann-Leitner ES, Angov E. Histidine affinity tags affect MSP1(42) structural stability and immunodominance in mice. Biotechnol J. 2012;7(1):133–47.

46 Wu J, Filutowicz M. Hexahistidine (His6)-tag dependent protein dimerization: a cautionary tale. Acta Biochim Pol. 1999;46(3):591–9.

47 Panek A, Pietrow O, Filipkowski P, Synowiecki J. Effects of the polyhistidine tag on kinetics and other properties of trehalose synthase from Deinococcus geothermalis. Acta Biochim Pol. 2013;60(2):163–6.

48 Buckner JD, Timpano KR, Zvolensky MJ, Sachs-Ericsson N, Schmidt NB. Implications of comorbid alcohol dependence among individuals with social anxiety disorder. Depress Anxiety. 2008;25(12):1028–37.

49 Buckner JD, Heimberg RG. Drinking behaviors in social situations account for alcohol-related problems among socially anxious individuals. Psychol Addict Behav. 2010;24(4):640–8.

50 Blumenthal H, Ham LS, Cloutier RM, Bacon AK, Douglas ME. Social anxiety, disengagement coping, and alcohol-use behaviors among adolescents. Anxiety Stress Coping. 2016;29(4):432–46.

51 Villarosa-Hurlocker MC, Madson MB. A latent profile analysis of social anxiety and alcohol use among college students. Addict Behav. 2020;104:106284.

52 Caruso MJ, Seemiller LR, Fetherston TB, Miller CN, Reiss DE, Cavigelli SA, et al. Adolescent social stress increases anxiety-like behavior and ethanol consumption in adult male and female C57BL/6J mice. Sci Rep. 2018;8(1):10040.

53 Simons JS, Simons RM, O’Brien C, Stoltenberg SF, Keith JA, Hudson JA. PTSD, alcohol dependence, and conduct problems: Distinct pathways via lability and disinhibition. Addict Behav. 2017;64:185–93.

54 Becker JB, Koob GF. Sex Differences in Animal Models: Focus on Addiction. Pharmacol Rev. 2016;68(2):242–63.

55 Paletta P, Bass N, Aspesi D, Choleris E. Sex Differences in Social Cognition. Curr Top Behav Neurosci. 2022.

56 Bredewold R, Veenema AH. Sex differences in the regulation of social and anxiety-related behaviors: insights from vasopressin and oxytocin brain systems. Curr Opin Neurobiol. 2018;49:132–40.

57 Li Y, Dulac C. Neural coding of sex-specific social information in the mouse brain. Curr Opin Neurobiol. 2018;53:120–30.

58 Knoedler JR, Inoue S, Bayless DW, Yang T, Tantry A, Davis CH, et al. A functional cellular framework for sex and estrous cycle-dependent gene expression and behavior. Cell. 2022;185(4):654–71 e22.

59 Kopachev N, Netser S, Wagner S. Sex-dependent features of social behavior differ between distinct laboratory mouse strains and their mixed offspring. iScience. 2022;25(2): 103735.

60 Verhagen M, van der Meij A, van Deurzen PA, Janzing JG, Arias-Vasquez A, Buitelaar JK, et al. Meta-analysis of the BDNF Val66Met polymorphism in major depressive disorder: effects of gender and ethnicity. Mol Psychiatry. 2010;15(3):260–71.

61 Jeanblanc J, He DY, Carnicella S, Kharazia V, Janak PH, Ron D. Endogenous BDNF in the dorsolateral striatum gates alcohol drinking. J Neurosci. 2009;29(43):13494–502.

62 Jeanblanc J, Logrip ML, Janak PH, Ron D. BDNF-mediated regulation of ethanol consumption requires the activation of the MAP kinase pathway and protein synthesis. Eur J Neurosci. 2013;37(4):607–12.

63 Moffat JJ, Sakhai SA, Ehinger Y, Phamluong K, Ron D. Brain derived neurotrophic factor in an orbitofrontal cortical-dorsolateral striatal circuit gates alcohol consumption bioRxiv. 2021:2021.09.10.459813.

64 Liu WZ, Zhang WH, Zheng ZH, Zou JX, Liu XX, Huang SH, et al. Identification of a prefrontal cortex-to-amygdala pathway for chronic stress-induced anxiety. Nat Commun. 2020;11(1):2221.

65 Qi CC, Wang QJ, Ma XZ, Chen HC, Gao LP, Yin J, et al. Interaction of basolateral amygdala, ventral hippocampus and medial prefrontal cortex regulates the consolidation and extinction of social fear. Behav Brain Funct. 2018;14(1):7.

66 Adhikari A, Topiwala MA, Gordon JA. Synchronized activity between the ventral hippocampus and the medial prefrontal cortex during anxiety. Neuron. 2010;65(2):257–69.

67 Okuyama T, Kitamura T, Roy DS, Itohara S, Tonegawa S. Ventral CA1 neurons store social memory. Science. 2016;353(6307):1536–41.

68 Sun Q, Li X, Li A, Zhang J, Ding Z, Gong H, et al. Ventral Hippocampal-Prefrontal Interaction Affects Social Behavior via Parvalbumin Positive Neurons in the Medial Prefrontal Cortex. iScience. 2020;23(3):100894.

69 Ewin SE, Morgan JW, Niere F, McMullen NP, Barth SH, Almonte AG, et al. Chronic Intermittent Ethanol Exposure Selectively Increases Synaptic Excitability in the Ventral Domain of the Rat Hippocampus. Neuroscience. 2019;398:144–57.

70 Marchant NJ, Campbell EJ, Whitaker LR, Harvey BK, Kaganovs.ky K, Adhikary S, et al. Role of Ventral Subiculum in Context-Induced Relapse to Alcohol Seeking after Punishment-Imposed Abstinence. J Neurosci. 2016;36(11):3281–94.

71 Massa SM, Yang T, Xie Y, Shi J, Bilgen M, Joyce JN, et al. Small molecule BDNF mimetics activate TrkB signaling and prevent neuronal degeneration in rodents. J Clin Invest. 2010;120(5):1774–85.

